# The RecQ DNA helicase Rqh1 promotes Rad3^ATR^ kinase signaling in the DNA replication checkpoint pathway of fission yeast

**DOI:** 10.1101/2020.04.10.036707

**Authors:** Nafees Ahamad, Saman Khan, Yong-jie Xu

## Abstract

Rad3 is the orthologue of ATR and the sensor kinase of the DNA replication checkpoint in *Schizosaccharomyces pombe*. Under replication stress, it initiates checkpoint signaling at the forks necessary for maintaining genome stability and cell survival. To better understand the checkpoint initiation process, we have carried out a genetic screen in fission yeast by random mutation of the genome looking for mutants with defects in Rad3 kinase signaling. In addition to the previously reported *tel2-C307Y* mutant (1), this screen has identified six mutations in *rqh1* encoding a RecQ DNA helicase. Surprisingly, these *rqh1* mutations except a start codon mutation are all in the helicase domain, indicating that the helicase activity of Rqh1 plays an important role in the replication checkpoint. In support of this notion, integration of two helicase-inactive mutations or deletion of *rqh1* generated a similar Rad3 signaling defect and heterologous expression of human RECQ1, BLM and RECQ4 restored the Rad3 signaling and partially rescued a *rqh1* helicase mutant. Therefore, the replication checkpoint function of Rqh1 is highly conserved and mutations in the helicase domain of these human enzymes may cause the checkpoint defect and contribute to the cancer predisposition syndromes.

## INTRODUCTION

Multiple cellular mechanisms ensure the accurate transmission of genetic material from one generation to the next. These include proper repair of DNA damage, protection of perturbed DNA replication forks, and the checkpoints that coordinate DNA replication and repair with cell cycle progression. Chromosomes are particularly vulnerable to DNA damage during replication as they are decondensed and single-stranded and highly accessible to damaging agents. In addition, other factors such as excessive trinucleotide repeats or insufficient supply of dNTPs can also perturb DNA replication. Therefore, DNA replication is subject to exquisite regulations. One such regulation mechanism is the DNA replication checkpoint (DRC). The DRC senses the problems and activates cellular responses such as increased production of dNTPs, cell cycle delay, fork stabilization, and suppression of late firing origins, which work in concert to minimize the mutation rate and ensure accurate and complete duplication of the genome before the cell division. Consistent with its importance in genome stability, the DRC is conserved from yeasts to humans, and defects in the cell signaling pathway cause a wide range of developmental and cancer predisposition syndromes. Although debatable, mutations generated by errors in DNA replication followed by mistakes in repair likely contribute significantly to sporadic cancers [see reviews in references (2-4)].

The current model envisages that a related set of sensor proteins in all eukaryotes that assemble at perturbed forks for the DRC signaling. Among the sensors, ATR is the protein kinase that works with its co-factor ATRIP (ATR-interacting protein) and the 9-1-1 (Rad9-Rad1-Hus1) complex to initiate the checkpoint signaling by phosphorylating various substrates including the mediators and effector kinase (5-7). The activated mediators channel the signal to the effector kinase (8,9). Once activated, the effector kinase diffuses away from the fork and relays the checkpoint signal to various cellular structures to stimulate the cellular responses mentioned above. However, the exact mechanism by which the checkpoint signaling is initiated at the perturbed forks remains incompletely understood (10-12).

To better understand the checkpoint initiation process (10,11), we have searched for new DRC mutants in *S. pombe* by random mutation of the genome looking for mutants with enhanced sensitivity to hydroxyurea (HU). HU perturbs DNA replication by inhibiting ribonucleotide reductase (RNR), a highly conserved enzyme that provides dNTPs for DNA replication and repair (13). HU slows down the polymerase movement at forks and consistent with this mechanism, HU resistance has been observed in cells overexpressing RNR or expressing a mutant RNR (14-16). Studies in various eukaryotic organisms have shown that the primary cellular response to HU treatment is the activation of the DRC (3,17). We have previously reported one of the screened mutants *tel2-C307Y* that is highly sensitive to the replication stress induced by HU or DNA damage (1). Here we report our characterization of a group of *rqh1* mutants.

Rqh1 is a member of the RecQ DNA helicase family that is conserved from bacteria to humans with an important role in genome surveillance. There are five RecQ helicases RECQ1, BLM, WRN, RECQ4 and RECQ5 in humans. Each of the five human helicases plays critical roles in maintaining genome stability. The importance of these enzymes in genome stability is underscored by the fact that mutations in all these helicases are linked to cancers or heritable cancer-prone syndromes such as Bloom’s, Werner’s and Rothmund-Thomoson syndromes [see reviews (18-20)]. Our genetic screen identified six mutations in *rqh1* from a group of nine newly screened *hus* mutants. Except a start codon mutation that depletes Rqh1, the rest five mutations are all in the helicase domain of Rqh1. One truncation mutant lacking most of the helicase domain behaves like *rqh1* null cells. The rest four mutations substitute the amino acids that are all highly conserved in RecQ DNA helicases and thus sensitize *S. pombe* to HU and DNA damage. Further studies on these mutants show that Rqh1 promotes Rad3 kinase signaling under replication stress and unlike the checkpoint mediator function of Sgs1 in *S. cerevisiae* (21,22), the DRC function of Rqh1 is likely mediated by the helicase activity in *S. pombe*.

## RESULTS

### Screening of *hus9* mutant with enhanced sensitivities to HU and DNA damage

We have carried out a large-scale *hus* (HU sensitive) screen in *S. pombe* looking for new DRC mutants and accumulated >1000 primary mutants. After backcrossing three times to remove bystander mutations and the crossing with DRC mutants to remove known mutations, a small set of new *hus* mutants was screened that likely causes DRC defects. Here, we first report the characterization of *hus9*, a representative of the linkage group of nine *hus* mutants and then the characterization of all *hus* mutants in this group. Preliminary results showed that phosphorylation of the DRC mediator Mrc1^Claspin^ by Rad3 (9,12) was significantly compromised in this mutant. We therefore decided to investigate on *hus9.*

We first examined the sensitivities of *hus9* to HU and DNA damage by using the spot assay described in Materials and Methods. In *S. pombe*, Rad3 is the master checkpoint kinase responsible for activation of both the DRC and the DNA damage checkpoint (DDC) pathways (23) (Fig. 1A). The ATM orthologue Tel1 contributes minimally to the checkpoints. Deletion of *rad3* sensitizes *S. pombe* to both HU and DNA damage due to a lack of the checkpoint functions. As shown in the top panels of Fig. 1B, while *S. pombe* lacking *rad3, mrc1* or *cds1* were highly sensitive to HU, the cells lacking *chk1*, the effector kinase of the DDC, was less sensitive, suggesting that the replication stress induced by HU is mainly dealt by the DRC. Under similar conditions, the *hus9* mutant was found sensitive to HU and the sensitivity was comparable to that of *mrc1* and *cds1*. We then tested the DNA damaging agents methylmethanesulfonate (MMS) and UV (Fig. 1B, middle panels). Unlike the HU treatment, *chk1* mutant was more sensitive to MMS and UV than *cds1*, indicating that the DNA damage that occurs at G2, the major cell cycle stage in *S. pombe*, is mainly dealt by the DDC. The *hus9* mutant was also sensitive to MMS and UV and the sensitivity was comparable to that of *rad3*. We then examined the sensitivity of *hus9* to bleomycin (Fig. 1B, bottom panels), an antibiotic that generates single-strand and double-strand breaks in chromosomal DNA (24). *hus9* was sensitive to bleomycin with a sensitivity even slightly higher than *rad3*.

**FIG 1.**
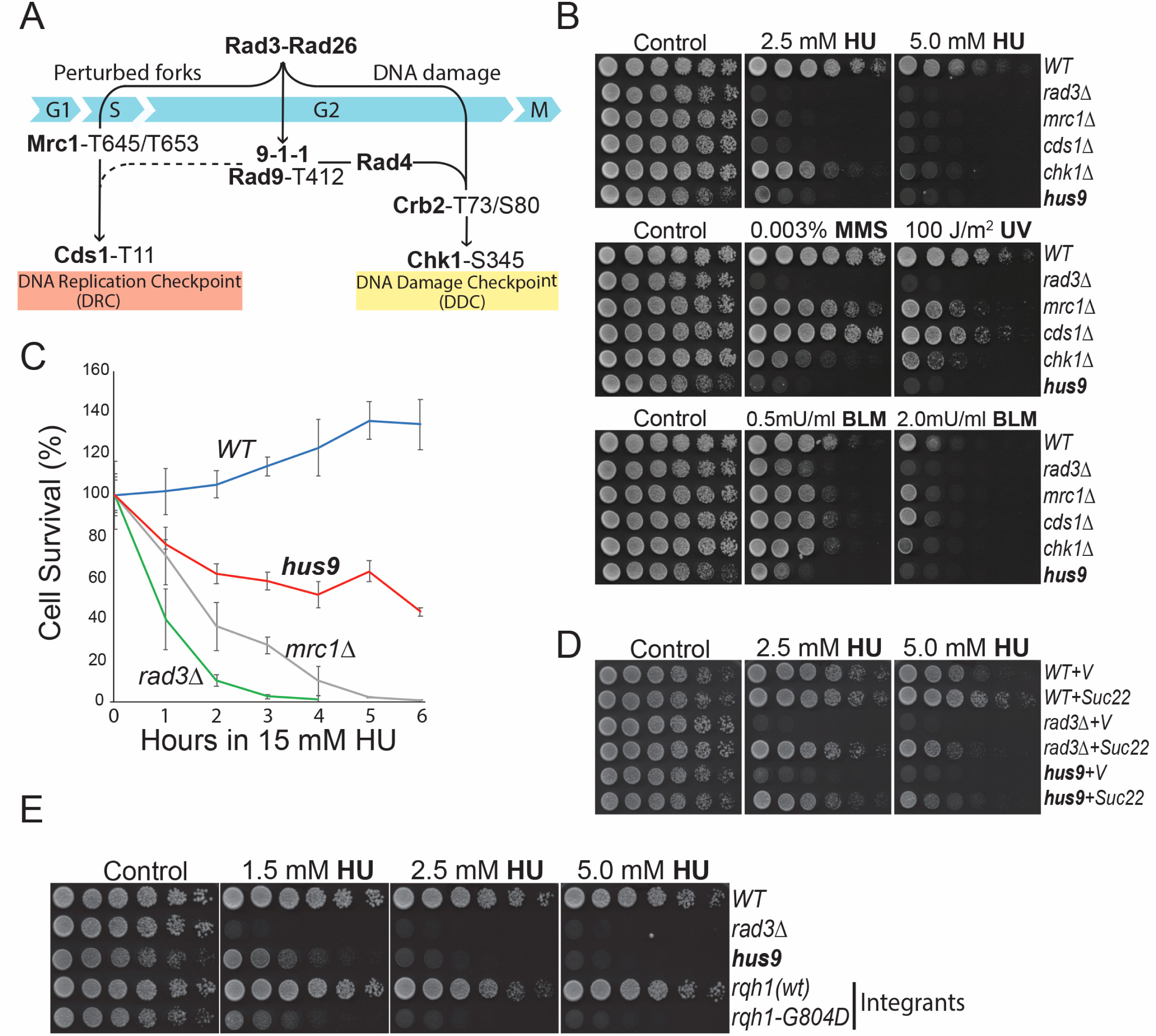
Sensitivity of the newly identified *hus9* mutant to HU and DNA damage. (**A**) Rad3 kinase signaling in the DRC (left) and the DDC (right) pathways of *S. pombe*. The serine and threonine residues phosphorylated by Rad3 are indicated. In the presence of perturbed replication, Rad3 phosphorylates Mrc1 and Cds1 to activate the DRC (9,12,43). When DNA damage occurs outside S phase or forks collapse, such as that in HU-treated *mrc1* or *cds1* cells, Rad3 phosphorylates Crb2^53BP1/Rad9^ and Chk1 to activate the DDC (46,48,49). Phosphorylation of Rad9 in the 9-1-1 complex is required for activation of both DRC and DDC (12,45). In the DDC pathway, phosphorylated Rad9 recruits Rad4^TopBP1/Dpb11^ to promote Crb2 and Chk1 activation. However, the function of phosphorylated Rad9 in promoting Cds1 activation remains unknown (dash line). (**B**) Sensitivities of *hus9* mutant (NF198) to HU, MMS, UV and bleomycin (BLM) were examined by the spot assay described in Materials and Methods. Wild type (TK48) and *rad3* (NR1826), *mrc1* (YJ15), *cds1* (GBY191) and *chk1* (TK197) mutants were used as controls. (**C**) Sensitivity of *hus9* to acute HU treatment in liquid medium. Cells used in B were incubated in YE6S containing 15 mM HU. At the indicated time points, cells were spread onto YE6S plates to recover for 3 days. Colonies were counted and presented in percentages. Error bars represent means and standard deviations (SD) of triplicates. (**D**) Overexpression of the RNR small subunit Suc22 rescues *hus9*. Suc22 was expressed in *S. pombe* on a vector under the control of its own promoter. V: vector. (**E**) The *hus* phenotype of *hus9* is caused by a single missense mutation in *rqh1*. The identified *rqh1-G804D* mutation was integrated at the genomic locus in wild type *S. pombe* (see Fig S2 for details). As control, wild type *rqh1* was integrated by the same method. HU sensitivity of the representative integrants of wild type (NF82) and mutant *rqh1* (NF83) was determined by spot assay. For comparison, *rad3* and *hus9* mutants were included.

We have recently screened a set of *hus* mutants of various metabolic pathways. These metabolic mutants such as *erg11-1* and *hem13-1* are highly sensitive to chronic exposure of HU as determined by the spot assay (25,26). However, they are not sensitive to DNA damage and the acute treatment of HU in liquid cultures. Upregulation of Suc22, the small subunit of RNR and the major regulatory target of DRC in *S. pombe* can rescue all DRC mutants but not the metabolic mutants. We then treated *hus9* with HU in liquid culture and found it was also sensitive and the sensitivity was less than *mrc1* and *rad3* (Fig. 1C). The higher HU sensitivity observed in the spot assay was likely due to oxidative stress (see Discussion). We also tested Suc22 and found its upregulation rescued *hus9* like in *rad3* (Fig. 1D). Tetrad dissection showed that crosses of *hus9* with all known DRC mutants generated spores with wild type HU resistance (Fig. S1), suggesting that *hus9* is not allelic to the checkpoint mutants and likely a new DRC mutant.

### *hus9* carries two missense mutations in *rqh1*

To identify the mutated gene, we transformed *hus9* with genomic DNA expression libraries marked with *ura4*. Colonies grown on medium lacking uracil were replicated onto plates containing HU to screen those with conferred HU resistance. Plasmids recovered from the yeast colonies were subjected to restriction enzyme digestions to remove those with *suc22*. Subsequent DNA sequencing identified *rqh1* and two G to A missense mutations at the *rqh1* genomic locus of *hus9*, which substitute Gly^767^ and Gly^804^ with Asp in the helicase domain of Rqh1 (Fig. S2A). Expression of wild type *rqh1* on a vector fully rescued *hus9* while the mutant *rqh1* did not (Fig. S2B, mid part). Separation of the two mutations showed that while *rqh1* containing the G767D mutation rescued *hus9, rqh1* containing G767D+G804D or G804D did not. To confirm, we integrated the mutations at the genomic locus in wild type *S. pombe* by using the method diagrammed in Fig. S2C. As the control, wild type *rqh1* was integrated by the same method. The integrants were screened by colony PCRs, backcrossed once to ensure single-copy integration, and then confirmed by PCRs (Fig. S2D) and Western blotting (Fig. S2E). The HU sensitivity of the resulting integrants was assessed by spot assay (Fig. S2B, lower part). The result showed that while the G767D integrant was resistant to HU like wild type cells, the integrants of *G804D* or G767D+G804D were sensitive like *hus9*. We also deleted *rqh1* by replacing the whole ORF with *ura4* and found that *hus9* showed a similar or slightly higher HU sensitivity (see below) as the *rqh1* deletion cells (Fig. S2B, upper part). Fig. 1E is a representative of the repeated spot assay, showing that a single G804D mutation in *rqh1* sensitizes *S. pombe* to HU and DNA damage. We hereafter renamed *hus9* as *rqh1-G804D* in the following studies.

*Rqh1* was originally identified in *S. pombe* as *hus2* (27,28) and *rad12* (29). Although studies in *S. cerevisiae* have shown that the Rqh1 homolog Sgs1 has checkpoint functions (21,22), earlier studies in fission yeast showed that Rqh1 mainly function in DNA repair, recombination, and fork recovery (27,29-34). Rqh1 is a member of the SF2 (superfamily 2) RecQ DNA helicase family that are highly conserved from bacteria to humans. There are five RecQ helicases in humans: RECQ1, BLM, WRN, RECQ4 and RECQ5 that play distinct and sometimes overlapping roles in maintaining genomic integrity through extensive involvement in DNA metabolism (18,19,35). Defects in BLM, WRN or RECQ4 cause the monogenic Bloom’s, Werner’s, and Rothmund-Thomson syndromes that are characterized by cancer predisposition or premature aging. Recent studies have also associated mutations of RECQL1 and RECQL5 to cancer predisposition or increased resistance to chemotherapies (36-40). All RecQ helicases share the ATP-dependent 3’-5’ DNA unwinding activity. Although *S. pombe* possess additional homologs (41,42), Rqh1 is generally believed to be the only RecQ helicase that functions at perturbed forks, which promotes unambiguous studies. In budding yeast, Sgs1 functions in the DRC as a mediator that recruits Rad53^CHK2/Cds1^ to be activated by Mec1^ATR/Rad3^ (21,22). Since our preliminary data suggest that Rqh1 promotes the phosphorylation of Mrc1 by Rad3, we decided to further investigate *rqh1-G804D* and its potential effect on DRC by using the set of phospho-specific antibodies that we have made available in the previous studies (9,12,43).

### HU arrests *rqh1-G804D* cells in S phase

As mentioned above, HU arrests the metabolic mutants at G2/M, not S phase, as they are killed by HU via the mechanisms unrelated to replication stress (25,26). We then examined the cell cycle progression by flow cytometry (Fig. 2A). In the presence of 15 mM HU, wild type cells were increasingly arrested at S phase in 1 to 3 h. Further incubation did not completely stop DNA synthesis and eventually finished the bulk of DNA synthesis in ∼7 to 8 h. *rad3* and *mrc1* cells were also arrested in S phase after 3 h in HU. However, these checkpoint mutants, and particularly *rad3* could not properly synthesize DNA in HU. Under similar conditions, HU arrested majority of *rqh1-G804D* at S phase although a small number of cells remained at G2/M after 3 to 4 h incubation. Like *rad3* and *mrc1, rqh1-G804D* also failed to continue DNA synthesis in HU, which suggests a defect in cellular response to HU.

**FIG 2.**
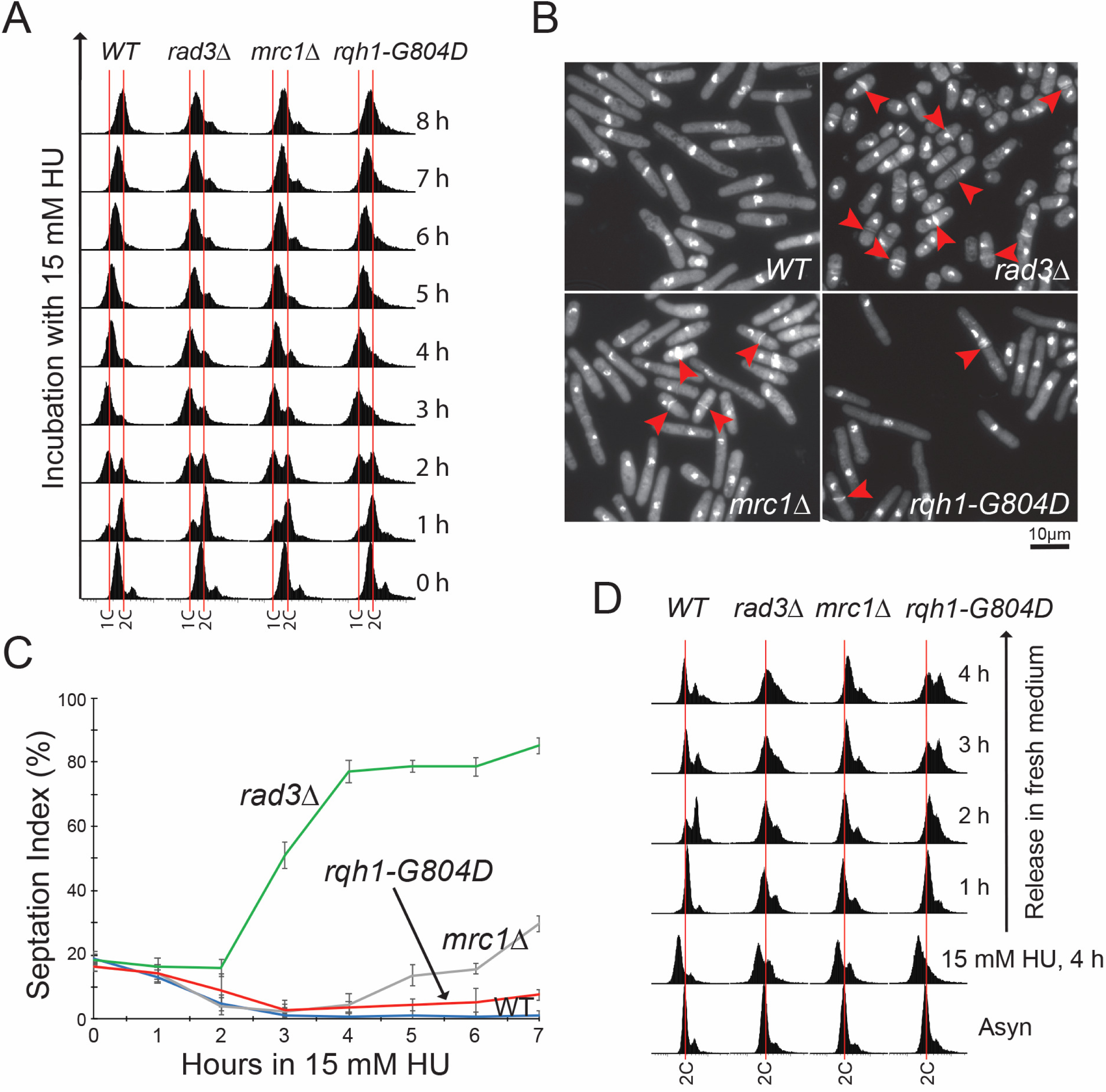
The DRC defect in *rqh1-G804D* mutant. (**A**) Cell cycle analysis of *rqh1-G804D*. Wild type, *rad3, mrc1* and *rqh1-G804D* cells used in Fig. 1C were incubated with 15 mM HU and analysed by flow cytometry every hour during the incubation. Red lines indicate 1C and 2C DNA contents. (**B**) *rqh1-G804D* cells elongated and showed the *cut* phenotype in HU. Cells were treated with HU for 6 h as in A, fixed onto glass slides by brief heating, stained with Hoechst and Blankophor for microscopic examination. Arrows indicate the *cut* cells. (**C**) A small number of *rqh1-G804D* cells undergo cell division in HU. Cells were incubated with HU as in A and fixed in 2.5% glutaraldehyde at the indicated time points. The fixed cells were washed and then stained as in B. ≥150 cells were counted under microscope for each sample, repeated three times. The cells with a septum are shown in percentages. (**D**) *rqh1-G804D* cells did not fully recover from HU arrest. The *S. pombe* strains used in A were arrested in HU for 4 h and then released in fresh medium. Cell cycle progression was monitored every hour during the release.

### Defects in recovery from replication stress

In the presence of HU, the DRC is activated to stimulate dNTP production and a delay in mitosis so that the cells can complete DNA synthesis before the division. The DRC mutants, however, proceed into mitosis, generating the *cut* phenotype with two daughters having unequal amounts or lacking genomic DNAs. We stained the cells with Hoechst for genomic DNAs and Blankophor for septum and examined cell septation during the HU treatment (Fig. 2B and C). After wild type cells were treated with HU for 3 h, cell division was almost completely suppressed and remained suppressed during the rest of HU treatment (Fig. 2C), suggesting an activated DRC. In contrast, *rad3* mutant showed a robust cell division activity in HU. As a result, most of the cells were short and showed the *cut* phenotype (Fig. 2B, arrows). The *mrc1* mutant initially slowed down cell division during the first 1 to 3 h. After 3 to 4 h, the cells began to divide, generating the *cut* cells (Fig. 2B and C). However, unlike the HU-treated *rad3* cells that were short due to the lack of both DRC and DDC, the *mrc1* cells elongated because the collapsed forks activate the DDC (9,44). The *rqh1-G804D* also showed the *cut* cells in HU and the septation index was lower than *mrc1* during the HU treatment (Fig. 2B and C). The smaller number of *cut* cells observed in *rqh1-G804D* is correlated with the milder HU sensitivity as compared with the *mrc1* mutant (Fig. 1C). Since the HU-treated *rqh1-G804D*, including the *cut* cells, were elongated, the DDC is likely activated as in *mrc1* cells (see below). Consistent with this result, the elongated and *cut* cells have also been described in HU-treated *S. pombe* lacking *rqh1* (27).

We then monitored cell recovery from HU by flow cytometry (Fig. 2D). After 4 h treatment in HU, majority of wild type and the mutant cells were arrested at S phase. When released in fresh medium, wild type cells fully recovered and returned to normal cell cycle in ≤ 4 h. In contrast, *rqh1-G804D* failed to fully recover like *rad3* and *mrc1* cells. These results strongly suggest a defect in DRC or cell recovery (27) from the S phase arrest by HU.

### Compromised Rad3 signaling in the DRC

Because our preliminary data showed that Rad3 phosphorylation of Mrc1 was significantly reduced in *rqh1-G804D*, we then examined the potential DRC defect by monitoring Rad3 phosphorylation of Rad9, Mrc1 and Cds1 using the phospho-specific antibodies (9,12,43). Under replication stress, Rad3 phosphorylates two functionally redundant residues Thr^645^ and Thr^653^ in Mrc1 and Thr^412^ in Rad9 of the 9-1-1 complex (Fig. 1A, left). Phosphorylation of Mrc1 and Rad9 facilitates the phosphorylation of Thr^11^ in Cds1^CHK2/Rad53^ by Rad3, which promotes the autophosphorylation of Cds1-Thr^328^ and autoactivation of the effector kinase (9,12,43). When DNA damage occurs outside S phase or forks collapse, such as that in HU-treated *mrc1* or *cds1* cells, Rad3 phosphorylates Rad9 and Crb2^53BP1/Rad9^ (45-47), which in turn facilitates the phosphorylation of Chk1-Ser^345^ by Rad3 (48,49), leading to the activation of the DDC (Fig. 1A, right).

We first examined Rad3-specific phosphorylation of Mrc1-Thr^645^, a representative of the two redundant sites in Mrc1 (Fig. 3A) (9). After 3 h HU treatment, while the phosphorylation was undetectable in *rad3*Δ, it significantly increased in wild type cells. Because the activated DRC upregulates Mrc1 (50,51), the level of Mrc1 in HU-treated *rad3* cells was lower than in wild type cells. Under similar conditions, Mrc1 phosphorylation in *rqh1-G804D* was reduced to 35% (±1.0%, n=3) of wild type level (Fig. 3A and S3A). We then treated the cells with 0.01% MMS for 90 min, similar reduction of Mrc1 phosphorylation was observed in *rqh1-G804D* (Fig 3B and S3B). To eliminate the cell cycle effect, we examined Mrc1 phosphorylation every hour during the HU treatment (Fig. S4A and B). While the phosphorylation was significantly increased in wild type cells during the first 3 h, it eventually decreased during the remaining time of the treatment, which is consistent with the flow cytometry data shown in Fig. 2A. In *rqh1-G804D*, Mrc1 phosphorylation was significantly lower at all time-points examined although it followed a similar kinetics as in wild type cells.

**FIG 3.**
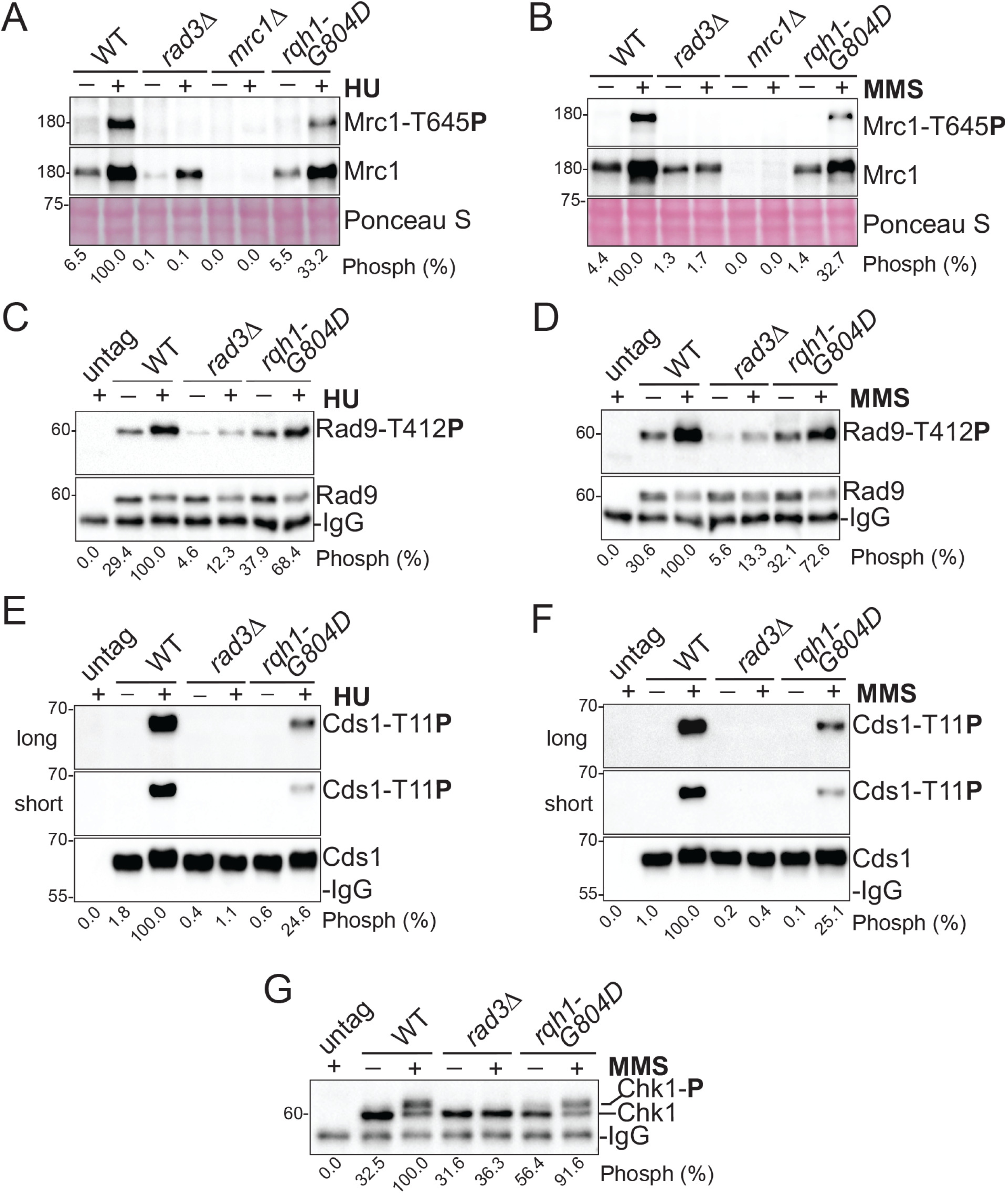
Compromised Rad3 kinase signaling in the DRC pathway. (**A**) HU-induced phosphorylation of Mrc1 by Rad3 was reduced in *rqh1-G804D*. Wild type and mutant strains used in Fig. 1B were treated with (+) or without (-) 15 mM HU for 3 h. Phosphorylation of Mrc1 (upper panel) was detected in whole cell lysates prepared by the TCA method using phospho-specific antibodies. The same blot was stripped and reprobed with anti-Mrc1 antibodies (mid panel). A section of the Ponceau S-stained membrane is shown (bottom panel). The phosphorylation bands were quantified, and band intensities compared to that in HU-treated wild type cells are shown at the bottom. (**B**) MMS-induced phosphorylation of Mrc1 was also reduced in *rqh1-G804D*. The cells were treated with 0.01% MMS for 90 min and then lysed for Western blotting analysis and subsequent quantification as in A. (**C**) HU-induced phosphorylation of Rad9 by Rad3 was moderately affected in *rqh1-G804D*. Wild type and the indicated mutant cells were treated with or without 15 mM HU for 3 h. Rad9-HA was IPed and separated by SDS-PAGE for Western blotting. The blot was first probed with anti-HA antibody (lower panel), stripped, and then reprobed with the phospho-specific antibody (upper panel). (**D**) Rad9 phosphorylation in the presence of MMS was also moderately affected in *rqh1-G804D*. The cells were treated with 0.01% MMS for 90 min and analysed by Western blotting as in C (**E**) Phosphorylation of Cds1 by Rad3 was significantly reduced in HU-treated *rqh1-G804D*. The cells were treated HU as in A. Cds1-HA was IPed and then analysed by Western blotting using anti-HA antibody (bottom panel). The same membrane was stripped and then blotted with phosphor-specific antibody (upper two panels). (**F**) MMS-induced phosphorylation of Cds1 was also significantly reduced in *rqh1-G804D*. The cells were treated with MMS as in B and D and analysed as in E. (**G**) MMS-induced phosphorylation of Chk1 by Rad3 was minimally affected in *rqh1-G804D*. Wild type, *rad3* and *rqh1-G804D* cells were treated with MMS as in B, D and F. Chk1-HA was IPed from an equal number of cells for Western blotting using anti-HA antibody. All experiments in this figure were repeated ≥ 3 times and the quantification results are shown in Fig. S3.

Under normal conditions, Rad9 is phosphorylated at a basal level mainly by Rad3 in *S. pombe* (12) (Fig. 3C and S3C). After HU treatment, the phosphorylation was significantly increased in wild type cells. In untreated *rqh1-G804D*, the basal phosphorylation remained detectable under or slightly higher. After HU treatment, however, the phosphorylation was also increased to 70.5% (±1.5%, n=3) of wild type level, suggesting that Rad9 phosphorylation was moderately affected. This result was confirmed by MMS treatment (Fig. 3D and S3D) as well as the time course study in HU (Fig. S4C and D). Finally, we examined Rad3 phosphorylation of Cds1 and found that the phosphorylation was significantly decreased to only 24.8% (±2.0%, n=3) of wild type level in HU (Fig. 3E and S3E) and 26.5% (±5.0%, n=3) in MMS (Fig. 3F and S3F). This result was also confirmed by the time course study in HU (Fig. S4E and F). These results show collectively that under replication stress, although Rad9 phosphorylation was moderately affected, the *rqh1-G804D* mutation significantly compromised Rad3 phosphorylation of Mrc1 and Cds1.

### Minimal reduction of Rad3 signaling in the DDC

Next, we examined the Rad3 signaling in the DDC pathway (Fig. 1A, right). As described above, in the presence of MMS, Rad9 phosphorylation by Rad3 was only moderately affected in *rqh1-G804D* (Fig. 3D and S3D). Under the similar conditions, Rad3 phosphorylation of Chk1 was minimally affected and remained at 90% (±1.5%, n=3) of wild type level (Fig. 3G and 3SG). This result was confirmed by the time course study (Fig. S4G and H), showing that although the DRC was significantly compromised, the DDC remained functional, which explains the cell elongation observed in *rqh1-G804D* (Fig. 2B). Though more severe, the similar defects in the DRC and the DDC have also been described in the previously reported *tel2-C307Y* mutant (1).

To further investigate the checkpoint defects, we crossed *rqh1-G804D* into checkpoint mutants and assessed the drug sensitivities of the resulting double mutants by spot assay (Fig. S5A and B). The result showed that while the *rad3*Δ*rqh1-G804D* double mutant was slightly more sensitive than the single mutants in the presence of HU or DNA damage, the sensitivity of the double mutants containing *mrc1*Δ or *cds1*Δ is comparable to that of *rqh1-G804D*, suggesting that Rqh1 plays an important role in the DRC. Interestingly, while the *chk1*Δ*rqh1-G804D* double mutant showed similar sensitivities to HU, UV, and MMS as in *rqh1-G804D*, the mutation suppressed the sensitivity of *chk1*Δ to bleomycin. We then made the *chk1*Δ*rqh1*Δ double mutant and found that it showed a similar sensitivity as in *rqh1*Δ in the presence of HU, MMS, UV and bleomycin (Fig. S5B). This result shows that the suppression of the bleomycin sensitivity in *chk1*Δ was allele-specific and is consistent with the multiple functions of Rqh1 in maintaining genome integrity (27,31,34). To provide further evidence for the DRC function, we examined the HU sensitivity of *mrc1*Δ*rqh1-G804D* in liquid culture and found that the double mutant showed a similar sensitivity as *mrc1*Δ although a slightly higher sensitivity was observed during the initial stage of HU treatment (Fig. S5C). This result is consistent with the DRC function of Rqh1.

### Association of Rqh1 with perturbed forks

RecQ DNA helicases bind to various fork DNA structures and interact with the replisome proteins [see review (19)], suggesting that Rqh1 associates with the forks to promote the DRC signalling in fission yeast. To investigate, we examined the potential interactions between Rqh1 and the replisome proteins Mrc1 and RPA. For this purpose, we tagged Mrc1 and Rpa1 with HA epitope and crossed into *S. pombe* with myc tagged Rqh1. As shown in Fig. 6A, when Rqh1 was IPed, Rpa1 was Co-IPed with Rqh1 (Fig. 4A and E). In the reciprocal Co-IP, Rqh1 was Co-IPed with Rpa1 (Fig. 4B and E). Since the Co-IPs under similar conditions did not detect any interactions between Rqh1 and Cds1 (see Fig. S9) and addition of ethidium bromide did not affect the interaction (52), Rqh1 may physically interact with RPA like the budding yeast Sgs1 (22). As mentioned above, Mrc1 is upregulated by HU treatment (Fig. 4C and D, inputs). Like Rpa1-Rqh1, Rqh1 and Mrc1 were found Co-IPed with each other (Fig. 4C, D and F). Furthermore, the interactions between Rqh1 and Rpa1 and Mrc1 were enhanced by the HU treatment (Fig. 4E and F), indicating that Rqh1 enriches at the HU-treated forks. Although the Co-IPs cannot clearly show whether the interactions of Rqh1 with RPA and Mrc1 is direct or indirect, nonetheless, Rqh1 is likely recruited to the perturbed forks via interactions with RPA and Mrc1 or binding to its DNA substrates associated with the forks.

**FIG 4.**
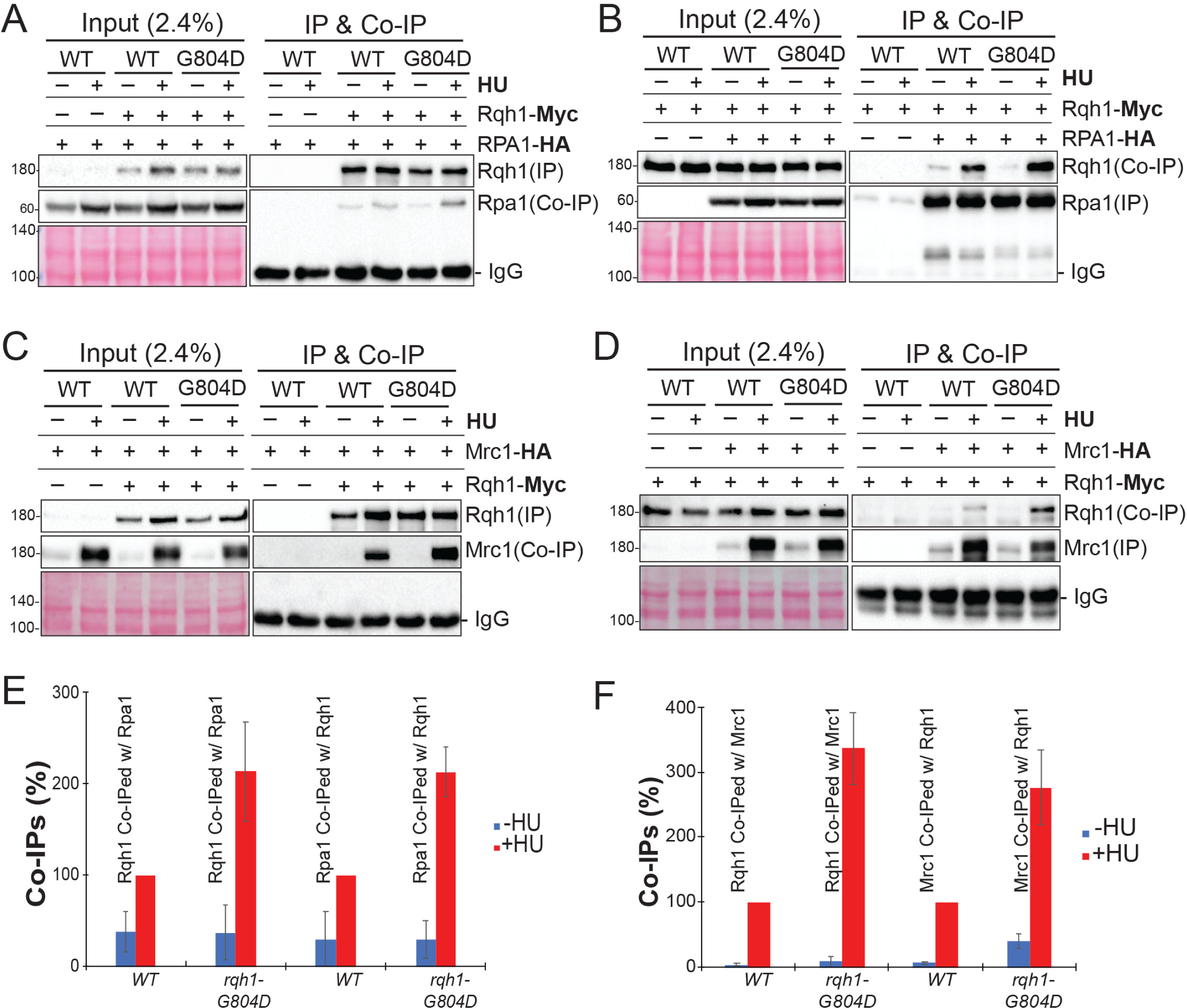
Interaction of Rqh1 with RPA and Mrc1. **(A)** Co-IP of Rpa1with Rqh1 was enhanced in a HU dependent manner. *S. pombe* with myc tagged wild type or mutant *rqh1* was crossed into cells expressing HA tagged Rpa1. Wild type (NF147) or mutant *rqh1* (NF164) cells were treated with (+) or without (-) 15 mM HU for 3 h. Rqh1 was IPed using anti-myc antibody (right upper panel) from cell extracts to detect Rpa1 Co-IPed with Rqh1 (lower right panel) as described in Materials and Methods. Untagged *rqh1* strain (YJ1586) was used as the control. 2.4% cells extracts were loaded on the same gel as the inputs (left three panels). A section of the Ponceau S-stained inputs is shown. (**B**) Rqh1 was Co-IPed with Rpa1. Rpa1 was IPed under similar conditions as in A with anti-HA antibody to detect Co-IPed Rqh1. (**C**) HU treatment enhanced the Co-IP of Mrc1 with Rqh1. The myc-*rqh1* strain was crossed into HA-*mrc1* strain as in A to generate wild type (NF157) and *rqh1-G804D* (NF168) strains. Rqh1 was IPed with anti-myc antibody as in A to detect Mrc1. (**D**) Reciprocal Co-IP of Rqh1 with Mrc1 was carried out in wild type and *rqh1-G804D* cells as in C. (**E**) & (**F**) Co-IPs shown in A, B, C and D were repeated ≥ 3 times. The Co-IPed bands were quantified and normalized with the inputs. After removing the non-specific bindings, intensities of the Co-IP bands were quantified and shown in percentages relative to HU-treated wild type cells in the presence (red columns) or absence (blue columns) of HU. Bars: means and SDs.

**FIG 5.**
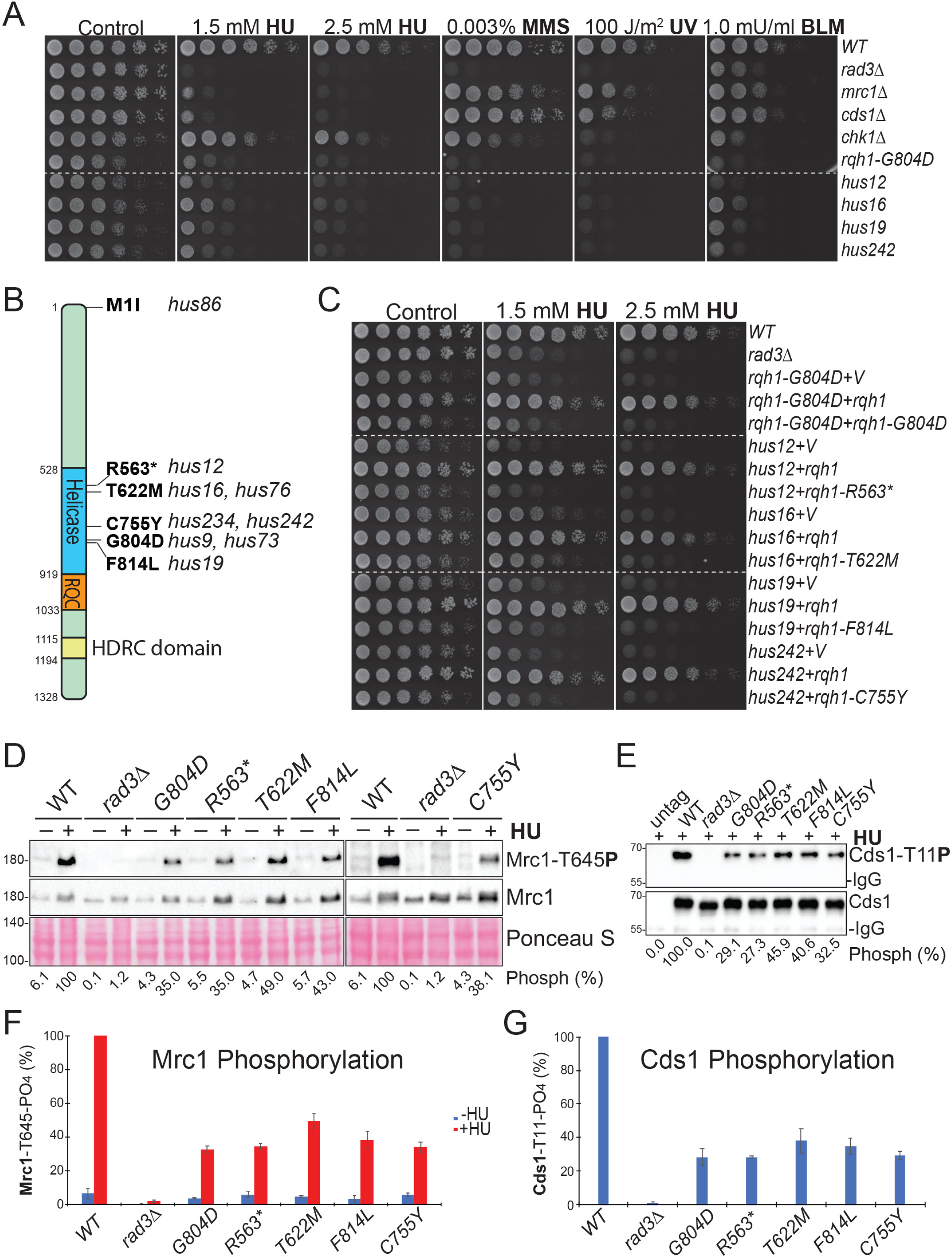
Drug sensitivity and the compromised DRC in other newly screened *rqh1* helicase mutants. (**A**) DNA sequencing of the nine newly screened *rqh1* mutants identified six uncharacterized mutations (see B). Sensitivities of the representative *rqh1* mutants to HU and the DNA damage were examined by spot assay. Dash line indicates discontinuity. (**B**) Schematics of Rqh1 with the conserved helicase, RQC, and HRDC domains. Relative locations of the mutations identified from the screened *hus* mutants are indicated. * in *hus12* indicates the truncation mutation. (**C**) Wild type and mutant *rqh1* were expressed in the indicated *rqh1* mutants on a vector under its own promoter. HU sensitivity was assessed by spot assay. (**D**) Rad3 phosphorylation of Mrc1 was compromised in all newly screened *rqh1* helicase mutants. Wild type and the representative mutants used in A were treated with or without 15 mM HU for 3 h. Mrc1 phosphorylation was detected in whole cell lysates using the phospho-specific antibodies and quantified as in Fig. 3A. (**E**) Rad3 phosphorylation of Cds1 was also compromised in the newly screened *rqh1* mutants. The cells used in D were treated with 15 mM HU for 3 h. Cds1-HA was IPed and then analysed by using the phosphor-specific antibody and anti-HA antibody as in Fig. 3E. (**F**) & (**G**) Experiments in D and E were repeated three times. The quantfictation results are shown for Mrc1 and Cds1, respectively. Bars: means and SDs.

**FIG 6.**
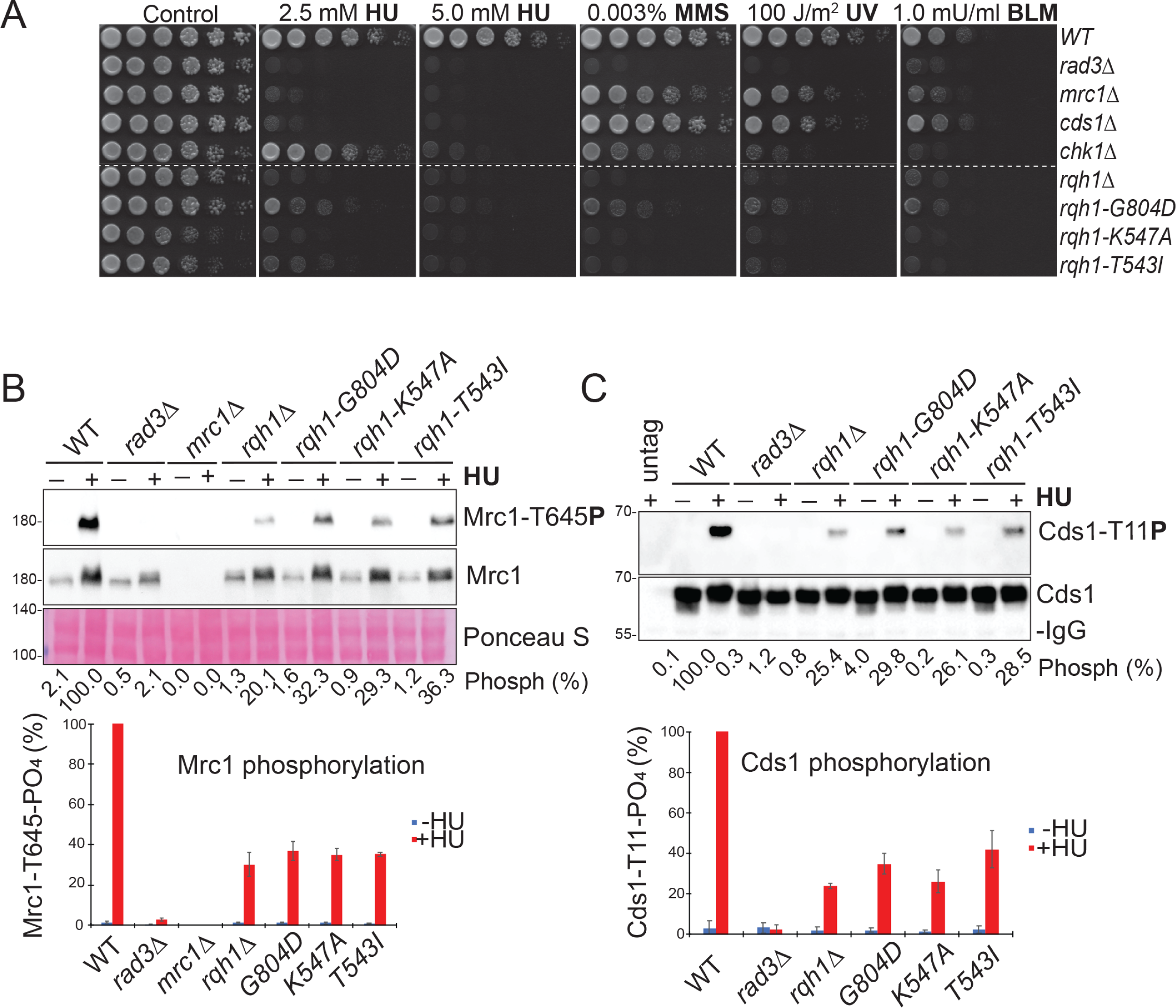
Deletion or helicase-inactive mutations of *rqh1* cause a similar Rad3 signaling defect as in *rqh1-G804D*. (**A**) Two helicase-inactive *rqh1-K547A* and *rqh1-T543I* mutation were integrated at the genomics as described for *rqh1-G804D* (Fig. S2C and S8A). Sensitivities of the indicated cells to HU and the DNA damaging agents were examined by spot assay. Wild type, the checkpoint mutants, and *S. pombe* lacking *rqh1* (NF121) were used as controls. Dash line indicates discontinuity. (**B**) Similar reduction of Rad3 phosphorylation was observed in *rqh1*Δ and the two helicase-inactive *K547A* and *T543I* mutants as in *rqh1-G804D*. Cells used in A were treated with or without 15 mM HU for 3 h before the analysis of Mrc1 phosphorylation. The experiment was repeated three times and the quantification results are shown in the bottom panel. Bars: means and SDs. (**C**) Reduced Rad3 phosphorylation of Cds1 in *rqh1*Δ and the helicase-inactive mutants. Cds1 phosphorylation was examined in cells carrying the indicated mutations treated with or without HU as in Fig 3. Quantification results from three independent experiments was shown in bottom panel.

In addition to Rqh1, fission yeast expresses additional RecQ helicase homologs (41,42), particularly the RecQ4 orthologue Hrq1 that plays important roles in DNA repair and genome stability. To see whether Hrq1 or other 3’-5’ helicases contribute to Rad3 signaling at the forks, we examined *S. pombe* lacking *hrq1, fml1 or fml2* (Fig. S6). The spot assay showed while the *rqh1-G804D* was highly sensitive to HU, *hrq1*Δ, *flm1*Δ and *fml2*Δ were not or minimally sensitive. Although the *hrq1*Δ and *fml1*Δ mutants were sensitive to bleomycin and MMS, respectively, the sensitivities were lower than in *rqh1-G804D*. Consistent with the drug resistance, Rad3 phosphorylation of Mrc1 in HU-treated *hrq1*Δ, *fml1*Δ and *fml2*Δ was at or near the wild type level (Fig. S6B and C), which excluded the possibility that these helicases function in the DRC.

### The DRC defect in other newly screened *rqh1* mutants

The helicase domain of Rqh1 is flanked by the long N- and C-terminal regions with less understood functions. In addition to *rqh1-G804D*, our *hus* screen has identified eight other mutants in the same linkage group that showed a similar HU sensitivity (Fig. 5A and B). To see whether mutations in the N- or C-terminal regions of *Rqh1* can also sensitize *S. pombe* to HU, we sequenced the *rqh1* genomic locus of these mutants and identified five more mutations (Fig. 5B). Among the mutations, *hus86* carries a start codon mutation, which likely affects the translation and our N-terminal tagging confirmed this notion (not shown). The remaining five mutations, to our surprise, are all located in the helicase domain. While *hus12* contains a truncation mutation that deletes the C-terminal 765 amino acids, the rest four mutations substitute the amino acids that are all highly conserved in RecQ DNA helicases (Fig. S7). To confirm the mutations, we expressed the mutant Rqh1 in their respective mutants on a vector under its own promoter (Fig. 5C). The results showed that while wild type Rqh1 fully rescued all mutants, the mutants expressing their mutant *rqh1* did not, which confirmed the mutations. Furthermore, we found that while *hus16* (T622M), *hus19* (F814L) and *hus242* (C755Y) were slightly less sensitive, the *hus12* (R563STOP) lacking the C-terminal half of Rqh1 was slightly more sensitive than *rqh1-G804D*, suggesting that the missense mutations did not fully abolish the function of Rqh1 (see below). We then examined the phosphorylation of Mrc1 (Fig. 5D and F) and Cds1 (Fig. 5E and G) in these mutants. The results showed clearly that in the presence of HU, all mutations significantly compromised the Rad3 phosphorylation of Mrc1 and Cds1 like in *rqh1-G804D*. Since these mutants were screened by random mutation of the genome, the helicase domain of Rqh1 must play an important role in promoting Rad3 signaling and cell survival in HU.

### The helicase-inactive *rqh1-G804D* mutation

All RecQ helicases have seven conserved motifs in their helicase domain (35) (Fig. S7). The mutated glycine in *rqh1-G804D* is highly conserved in motif V. Earlier studies have shown that mutation of this residue in other SF2 helicases eliminated the helicase and NTP hydrolysis activities (53,54), suggesting that the mutation abolishes the helicase activity of Rqh1. To investigate, we integrated two previously reported helicase-inactive T543I and K547A mutations (34) at the genomic locus. Western blotting showed that they were expressed at the same level as in wild type and *rqh1-G804D* cells (Fig. S8A). We found that in the presence of HU, MMS, UV and bleomycin, although slightly less sensitive, the *rqh1-G804D* behaved like the two helicase-inactive mutants and *S. pombe* lacking *rqh1* (Fig 6A). We then compared the phosphorylation of Mrc1 (Fig. 6B) and Cds1 (Fig. 6C) in these mutants. As expected, the phosphorylation was similarly reduced in *rqh1*Δ and the two helicase-inactive mutants as in *rqh1-G804D*, suggesting that although not completely, the newly screened mutations including *rqh1-G804D* eliminated most of the helicase activity of Rqh1. Previous studies showed that RecQ helicases may function as oligomers (55-57). If *rqh1-G804D* retains a functional helicase domain, co-expression of *rqh1-G804D* may rescue the helicase-inactive mutants. We then expressed *rqh1-G804D* in K547A and T543I and assessed the HU sensitivity. The result showed that *rqh1-G804D* did not have any rescuing effect (Fig. S8B). These results show that *rqh1-G804D* as well as the other newly identified *rqh1* mutations likely eliminate most of the helicase activity of Rqh1 and thus compromise the Rad3 signaling.

### Human BLM, RECQ1 and RECQ4 restore Rad3 signaling and rescue *rqh1-G804D*

*S. pombe* is an established model for studying the cellular mechanisms that are conserved in higher eukaryotes (58) and cross-species complementation of yeast mutations by human genes has been successfully utilized to elucidate the functional homology between human and yeast proteins (59,60). We then tested whether expression of the mammalian enzymes can rescue *rqh1-G804D*. To this end, human BLM, RECQ1, RECQ4, RECQ5 and mouse WRN were expressed from a vector under the control of *nmt1* promoter in *rqh1-G804D* (Fig. 7A). The spot assay showed that while WRN or RECQ5 did not shown any rescuing effect, BLM, RECQ1 and RECQ4 partially rescued *rqh1-G804D* in the presence of HU, UV and MMS. Western blotting confirmed the thiamine-regulated expression of all mammalian proteins in *rqh1-G804D* (Fig 7B). We then examined Rad3 phosphorylation of Mrc1 and found that consistent with the rescuing effect, the phosphorylation was significantly restored in *rqh1-G804D* expressing RECQ1, BLM, or RECQ4. Consistent with our data, an earlier study showed that while expression of BLM suppressed the HU sensitivity of a budding yeast *sgs1* mutant, WRN did not (61). Importantly, since human RECQ1 is the smallest homologue and contains mainly the helicase domain, its rescuing effect observed in *rqh1-G804D* strongly support our notion that it is the helicase activity of Rqh1 not the protein per se that plays an important role in promoting Rad3 signaling at the perturbed forks in *S. pombe*.

**FIG 7.**
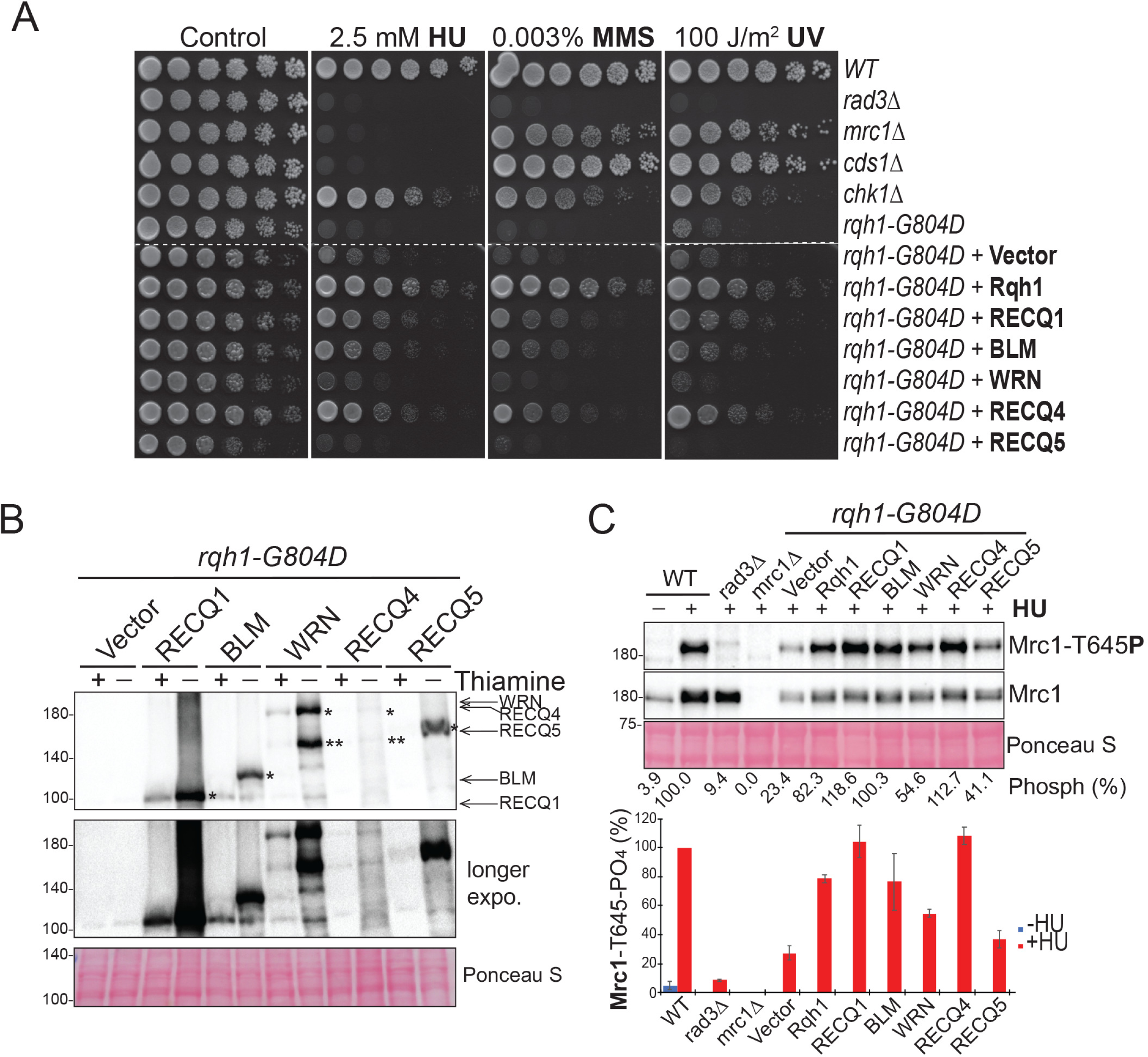
Complementation of *rqh1-G804D* by human RecQ DNA helicases. (**A**) Heterologous expression of human RECQ1, BLM or RECQ4, not human RECQ5 or mouse WRN rescued *rqh1-G804D*. The mammalian helicases were expressed in *rqh1-G804D* from a vector under *nmt1* promotor. The sensitivities of *rqh1-G804D* expressing the human enzymes to HU, MMS and UV were examined by spot assay. Wild type, the checkpoint mutants, *rqh1-G804D* and *rqh1-G804D* carrying an empty vector were used as the controls. (**B**) Thiamine-regulated expression of the mammalian helicases in *rqh1-G804D*. The myc-tagged mammalian helicases were expressed under the *nmt1* promotor. The expression levels were analysed in *rqh1-G804D* cultured in the presence (+) or absence (-) thiamine by Western blotting using anti-myc antibodies. A section of Ponceau S-stained membrane is shown as the loading control (bottom panel). While * indicates the full-length enzymes, ** indicates the degradation products. (**C**) Rad3 phosphorylation of Mrc1 was restored in *rqh1-G804D* expressing human RECQ1, BLM or RECQ4. Cells used in A were treated with 15 mM HU for 3 h. Phosphorylation of Mrc1 (upper panel) and Mrc1 (mid panel) were sequentially detected by using phospho-specific antibodies and the anti-Mrc1 antibodies as described in Fig. 3A. Quantification results from three repeated experiments are shown on the bottom. Bars: means and SDs.

## DISCUSSION

Our genetic screen has identified a new set of *hus* mutants with DRC defects. Characterization of nine *hus* mutants in the same linkage group identified six mutations in *rqh1* (Fig. 5B). Except the start codon mutation in *hus86*, all rest five mutations are in the helicase domain. One of the mutant carries a truncation mutation that behaves like *rqh1* null cells under all tested conditions. The rest four mutations substitute the amino acids that are all highly conserved in the RecQ DNA helicases (Fig. S7). Although subtle differences are observed in drug sensitivity and the DRC signalling assays, these four missense mutants behave quite similar. Consistent with their slightly milder HU sensitivity, the defect in Rad3 kinase signaling was slightly less severe than in *rqh1* null cells. These results, together with experiments on *rqh1-G804D* show that Rqh1 promotes Rad3 kinase signalling at the perturbed forks in *S. pombe*. At the molecular level, all newly screened mutations compromised Rad3 phosphorylation of Mrc1 and Cds1 in the presence of HU or MMS. At the cellular level, HU generates *cut* cells and the number of *cut* cells corroborates with the cell killing effect of HU in *rqh1-G804D*, the presentative of the newly screened nine *hus* mutants. We also found that the DDC is minimally affected in *rqh1-G804D*, which is consistent with the cell elongation and the compromised DRC observed in this mutant. Although our data provide an explanation for the sensitivity of *rqh1* mutants to replication stress induced by HU or DNA damage, they do not exclude the involvement of Rqh1 in fork restart and recombination repair (27,31,34,62). Although *S. cerevisiae* Sgs1 is known to function in the DRC (21,22), whether a similar function is conserved in *S. pombe* Rqh1 has been controversial (29,31,62). It is possible that the Cds1 kinase assays used in the previous studies are semi-quantitative that prevent accurate measurement of the partial DRC defect that we observed in the *rqh1* mutants. It is also possible that the difference is caused by allele-specific effects.

Comparing with *mrc1* null cells lacking the DRC, there is a difference between the HU sensitivity of *rqh1-G804D* as determined by spot assay (Fig 1B) and that in liquid cultures (Fig 1C). As mentioned above, this difference has been observed in the metabolic *hus* mutants that are killed by HU via oxidative stress or other mechanisms (25,26). It has also been observed in our previous study on several mutations of Rad4^TOPBP1/Dpb11^, a protein that functions in both replication initiation and the DDC (10). It is likely that the oxidative stress generated by perturbed DNA replication in *S. pombe* (63) may exacerbate the cell-killing effect of HU, particularly the chronic exposure as in spot assay (64). Rqh1, though not essential for cell growth, may contribute to normal DNA replication and forks stability, particularly at the hard-to-replicate chromosomal regions. In support of this possibility, a slightly increased Rad9 and Chk1 phosphorylation (Fig 3D and G) and longer G2/M delay (Fig 2 A and D) have been observed in *rqh1-G804D* under normal conditions. Therefore, *rqh1* mutations may minimally perturb normal DNA replication, which generate oxidative stress and thus enhance the chronic cell-killing effects of HU.

In *S. cerevisiae*, Sgs1 is phosphorylated by Mec1^ATR/Rad3^ to recruit Rad53^CHK2/Cds1^ as a checkpoint mediator for Rad53 activation (21,22,65). Our data show that Rqh1 likely promotes Rad3 kinase signalling at the forks by its helicase activity, not as a mediator in fission yeast. First, except the mutations truncating or depleting Rqh1, the missense mutations identified by our random genome-wide screen are all in the helicase domain, not the other parts of the protein. Second, the DRC is similarly compromised in *rqh1*Δ, the two helicase-inactive mutants as well as the newly screened *rqh1* mutants. Third, *cut* cells, a typical phenotype associated with DRC mutants, are observed in *rqh1-D804D* (Fig 2B and C) as well as in *rqh1*Δ cells (27). Fourth, unlike Sgs1, Rqh1 lacks the consensus docking sites for Cds1 via the phospho-peptide binding FHA domain of Cds1 (9,66). Consistent with this, our Co-IPs did not detect the interaction of Rqh1 with Cds1 (Fig S9) although the interaction of Sgs1 with Rad53 (22,65) and that of Rqh1 with Rpa1 and Mrc1 can be detected by the same method (Fig 4). Finally, heterologous expression of human RecQ helicases BLM, RECQ4, and particularly the smallest RecQ helicase RECQ1 that mainly contains the helicase domain rescues and restores Rad3 signaling in *rqh1-G804D*.

In addition to the DRC function of budding yeast Sgs1 mentioned above (21,22), RecQ, the founding member of the RecQ helicase family, initiates the SOS response at perturbed forks by its helicase activity in bacteria (67). The DRC functions have also been described for human RECQ1 (68), RECQ4 (69,70) and WRN (71-73). Although not directly involved in the DRC signaling, BLM is regulated by ATR phosphorylation for cell recovery from S-phase arrest (74). It is likely that the DRC function of Rqh1 is conserved from bacteria and yeasts to humans and the defects associated with the helicase mutations in the human enzymes contribute to the disease syndromes mentioned above. Future studies are needed to investigate the enzymatic activities of Rqh1 that promote the Rad3 signaling and the possible redundant factor(s) responsible for the remaining 20-30% Mrc1 phosphorylation in fission yeast. Furthermore, our humanized *S. pombe* may also provide an *in vivo* platform to understand the DRC defects and to screen for specific inhibitors of the human RecQ DNA helicases.

## MATERIALS AND METHODS

### Yeast strains and plasmids

The *S. pombe* strains were cultured at 30°C in YE6S (0.5% yeast extract, 3% dextrose and 6 supplements) or in EMM[6S] medium lacking an appropriate supplement following standard methods (75). Yeast strains, plasmids, and PCR primers used in this study are listed in Supplementary Table S1, S2, and S3, respectively. Mammalian RecQ helicases RECQL, pTRIP-CMV-puro-2A-BLM, pCMV-WS-Myc-WRN, and TAL2220 was obtained from Addgene (Cat #38890, #127641, #19273, and #36679, respectively). RECQ5 plasmid was kindly provided by Dr. Nicola Burgess-Brown at University of Oxford. These mammalian RecQ helicases was amplified by Phusion polymerase and cloned between NotI and XmaI sites in a yeast expression vector under the control of *nmt1* promotor. Cloned genes and the mutations were confirmed by DNA sequencing (Retrogen, San Diego).

### The *hus* screen

The method for screening new *hus* mutants has been described in our previous study (1).

### Integration of *rqh1* mutations

The *rqh1* expression cassette was cloned into the pIRT-2U vector between SacI and XmaI sites. An myc epitope and a *kan*^*r*^ marker were sequentially inserted in frame at the N-terminus of *rqh1* and after the terminator, respectively (see Fig. S2C). After digestion with NheI and PciI, a 7131 bp integration fragment was gel purified and transformed into the wild type TK7 strain. After an overnight recovery, cells were replicated onto plates containing 100 µg/ml G418 (Sigma). Colonies were screened by colony PCRs to confirm correct integrations at both 5’ and 3’ ends. The integrants were backcrossed once to ensure single copy integration and confirmed again by PCRs that cover the whole regions of the integration locus as well as Western blottings with anti-myc antibody as shown in Fig S2D and E.

### Drug sensitivity

Sensitivities to HU and various DNA damaging agents were determined by spot assay or in liquid medium as described in our previous studies (9,12,25,26). For the spot assay, 2 x 10^7^ cells/ml of logarithmically growing *S. pombe* were diluted in five-fold steps and spotted in 3 µl onto YE6S plates or YE6S plates containing the drugs at indicated concentrations. The plates spotted with the cells were dried before the treatment with UV (Stratalinker 2400). The plates were incubated at 30°C for 3 days and then photographed. All spot assays were repeated at least once. For acute HU sensitivity (28), liquid cell cultures were incubated with 15 mM HU. Every hour during the treatment, an equal number of cells were removed, diluted 1000-fold, spread onto three YE6S plates, and incubated at 30°C for 3 days for cell recovery. Colonies were counted and presented as percentages of the untreated cells.

### IP and Co-IP

1 × 10^8^ logarithmically growing cells were harvested and saved at −20°C in a 1.5 ml screw cap tube. The frozen cell pellets were lysed by mini-bead beater in the buffer containing 25 mM HEPES/NaOH (pH7.5), 50 mM NaF, 1 mM NaVO_4_, 10 mM NaP_2_O_7_, 40 mM ß-glycerophophate, 0.1% Tween 20, 0.5% NP-40, and protease inhibitors. The lysates were centrifuged at 16,000 g, 4°C for 5 min to clarify. Anti-HA or anti-myc antibody agarose resins (Santa Cruz) were prewashed 3x with Tris-buffered saline containing 0.05% Tween 20 (TBS-T) and incubated with 5% BSA in TBS-T for ≥ 30 min at 4°C. The cell extract was treated with 100 μg/ml ethidium bromide (Sigma) (52) for 1 h before incubation with the prewashed antibody resins by rotating in 2 ml tubes at 4°C for 2 h. The resins were washed 3x with TBS-T at 4°C for 20 min and then separated by SDS-PAGE followed by Western blottings with mouse monoclonal anti-HA (12CA5, Sigma) or anti-myc (9E10, Thermo Scientific) antibodies.

### Western blotting

The method for examining Rad3 phosphorylation of Mrc1-Thr^645^, Rad9-Thr^412^ and Cds1-Thr^11^ using the phosphor-specific antibodies has been described in our previously studies (1,12,43). The myc or HA tagged proteins was examined by Western blotting by using mouse monoclonal antibodies. For the Western analyses, 1 x 10^8^ logarithmically growing cells were fixed in 15% trichloroacetic acid on ice for ≥ 3 h and then lysed by mini-bead beater. The lysates from 2 - 4 x 10^6^ cells were separated by SDS-PAGE before transferring to the nitrocellulose membrane and stained with Ponceau S as the loading control (76). The blotting signal was detected by electrochemiluminescence using ChemiDoc XRS Imaging system (BioRad). Intensities of the specific bands were quantified and analysed by ImageLab (BioRad).

### Flow cytometry

1 ×10^7^ logarithmically growing cells were collected, fixed in ice-cold 70% ethanol, and then analysed by Accuri C6 flow cytometer as described in our previous studies (25,26).

### Microscopy

The cells were fixed directly onto uncoated glass slides by heating briefly at 75°C for 30 sec or in medium containing 2.5% glutaraldehyde at 4°C for ≥ 3 h. The glutaraldehyde-fixed cells were washed with PBS by centrifugation at 2300 g for 30 sec, stained in the same buffer with 5 µg/ml Hoechst33258 (Sigma-Aldrich) and 1:100 dilution of the Blankophor working solution (MP Biochemicals). The stained cells were examined using an Olympus EX41 fluorescent microscope. Images were captured with an IQCAM camera (Fast1394) using Qcapture Pro 6.0 software. ∼150 cells were counted under microscope for each sample and repeated three times. Images were also extracted into Photoshop (Adobe) to generate Fig. 2B.

## ACKNOWLEDGEMENT

We thank NBRP/YGRC in Japan, Drs Tom Kelly, Paul Russell, Michael Boddy and Matthew Whitby for sharing the yeast strains and Dr Nicola Burgess-Brown for sharing the RECQ5 plasmid. We also thank other members of the Xu lab for their help and support. This work was supported by NIH RO1 grant GM110132 to YJX.

## ABBREVATIONS

ATM: ataxia telangiectasia mutated;
ATR: ataxia telangiectasia and Rad3 related;
BLM: bleomycin;
*cut*: cell untimely torn;
DDC: DNA damage checkpoint;
DRC: DNA replication checkpoint;
HU: hydroxyurea;
*hus*: hydroxyurea sensitive;
IP: immunoprecipitation;
MMS: methylmethanesulfonate;
RNR: ribonucleotide reductase;
TCA: trichloroacetic acid;
UV: ultraviolet.

## REFERENCES

1. Xu, Y.J., Khan, S., Didier, A.C., Wozniak, M., Liu, Y., Singh, A. and Nakamura, T.M. (2019) A tel2 Mutation That Destabilizes the Tel2-Tti1-Tti2 Complex Eliminates Rad3(ATR) Kinase Signaling in the DNA Replication Checkpoint and Leads to Telomere Shortening in Fission Yeast. Mol Cell Biol, 39, e00175–00119.

2. Yazinski, S.A. and Zou, L. (2016) Functions, Regulation, and Therapeutic Implications of the ATR Checkpoint Pathway. Annu Rev Genet, 50, 155–173.

3. Iyer, D.R. and Rhind, N. (2017) The Intra-S Checkpoint Responses to DNA Damage. Genes (Basel), 8.

4. Golemis, E.A., Scheet, P., Beck, T.N., Scolnick, E.M., Hunter, D.J., Hawk, E. and Hopkins, N. (2018) Molecular mechanisms of the preventable causes of cancer in the United States. Genes Dev, 32, 868–902.

5. Zou, L. and Elledge, S.J. (2003) Sensing DNA damage through ATRIP recognition of RPA-ssDNA complexes. Science, 300, 1542–1548.

6. Kumagai, A., Lee, J., Yoo, H.Y. and Dunphy, W.G. (2006) TopBP1 activates the ATR-ATRIP complex. Cell, 124, 943–955.

7. Lee, J., Kumagai, A. and Dunphy, W.G. (2007) The Rad9-Hus1-Rad1 checkpoint clamp regulates interaction of TopBP1 with ATR. J Biol Chem, 282, 28036–28044.

8. Tanaka, K. and Russell, P. (2001) Mrc1 channels the DNA replication arrest signal to checkpoint kinase Cds1. Nat Cell Biol, 3, 966–972.

9. Xu, Y.J., Davenport, M. and Kelly, T.J. (2006) Two-stage mechanism for activation of the DNA replication checkpoint kinase Cds1 in fission yeast. Genes Dev, 20, 990–1003.

10. Yue, M., Zeng, L., Singh, A. and Xu, Y.J. (2014) Rad4 mainly functions in Chk1-mediated DNA damage checkpoint pathway as a scaffold protein in the fission yeast Schizosaccharomyces pombe. PLoS One, 9, e92936.

11. Bandhu, A., Kang, J., Fukunaga, K., Goto, G. and Sugimoto, K. (2014) Ddc2 Mediates Mec1 Activation through a Ddc1-or Dpb11-Independent Mechanism. PLoS Genet, 10, e1004136.

12. Yue, M., Singh, A., Wang, Z. and Xu, Y.J. (2011) The phosphorylation network for efficient activation of the DNA replication checkpoint in fission yeast. J Biol Chem, 286, 22864–22874.

13. Nordlund, P. and Reichard, P. (2006) Ribonucleotide reductases. Annu Rev Biochem, 75, 681–706.

14. Sneeden, J.L. and Loeb, L.A. (2004) Mutations in the R2 subunit of ribonucleotide reductase that confer resistance to hydroxyurea. J Biol Chem, 279, 40723–40728.

15. Choy, B.K., McClarty, G.A., Chan, A.K., Thelander, L. and Wright, J.A. (1988) Molecular mechanisms of drug resistance involving ribonucleotide reductase: hydroxyurea resistance in a series of clonally related mouse cell lines selected in the presence of increasing drug concentrations. Cancer Res, 48, 2029–2035.

16. Akerblom, L., Ehrenberg, A., Graslund, A., Lankinen, H., Reichard, P. and Thelander, L. (1981) Overproduction of the free radical of ribonucleotide reductase in hydroxyurea-resistant mouse fibroblast 3T6 cells. Proc Natl Acad Sci U S A, 78, 2159–2163.

17. Elledge, S.J., Zhou, Z. and Allen, J.B. (1992) Ribonucleotide reductase: regulation, regulation, regulation. Trends Biochem Sci, 17, 119–123.

18. Larsen, N.B. and Hickson, I.D. (2013) RecQ Helicases: Conserved Guardians of Genomic Integrity. Adv Exp Med Biol, 767, 161–184.

19. Croteau, D.L., Popuri, V., Opresko, P.L. and Bohr, V.A. (2014) Human RecQ helicases in DNA repair, recombination, and replication. Annu Rev Biochem, 83, 519–552.

20. Urban, V., Dobrovolna, J. and Janscak, P. (2017) Distinct functions of human RecQ helicases during DNA replication. Biophys Chem, 225, 20–26.

21. Frei, C. and Gasser, S.M. (2000) The yeast Sgs1p helicase acts upstream of Rad53p in the DNA replication checkpoint and colocalizes with Rad53p in S-phase-specific foci. Genes Dev, 14, 81–96.

22. Hegnauer, A.M., Hustedt, N., Shimada, K., Pike, B.L., Vogel, M., Amsler, P., Rubin, S.M., van Leeuwen, F., Guenole, A., van Attikum, H. et al. (2012) An N-terminal acidic region of Sgs1 interacts with Rpa70 and recruits Rad53 kinase to stalled forks. EMBO J, 31, 3768–3783.

23. Bentley, N.J., Holtzman, D.A., Flaggs, G., Keegan, K.S., DeMaggio, A., Ford, J.C., Hoekstra, M. and Carr, A.M. (1996) The Schizosaccharomyces pombe rad3 checkpoint gene. EMBO J, 15, 6641–6651.

24. Povirk, L.F. (1996) DNA damage and mutagenesis by radiomimetic DNA-cleaving agents: bleomycin, neocarzinostatin and other enediynes. Mutat Res, 355, 71–89.

25. Xu, Y.J., Singh, A. and Alter, G.M. (2016) Hydroxyurea induces cytokinesis arrest in cells expressing a mutated sterol-14alpha-demethylase in the ergosterol biosynthesis pathway. Genetics, 204, 959–973.

26. Singh, A. and Xu, Y.J. (2017) Heme deficiency sensitizes yeast cells to oxidative stress induced by hydroxyurea. J Biol Chem, 292, 9088–9103.

27. Stewart, E., Chapman, C.R., Al-Khodairy, F., Carr, A.M. and Enoch, T. (1997) rqh1+, a fission yeast gene related to the Bloom’s and Werner’s syndrome genes, is required for reversible S phase arrest. Embo J, 16, 2682–2692.

28. Enoch, T., Carr, A.M. and Nurse, P. (1992) Fission yeast genes involved in coupling mitosis to completion of DNA replication. Genes Dev, 6, 2035–2046.

29. Davey, S., Han, C.S., Ramer, S.A., Klassen, J.C., Jacobson, A., Eisenberger, A., Hopkins, K.M., Lieberman, H.B. and Freyer, G.A. (1998) Fission yeast rad12+ regulates cell cycle checkpoint control and is homologous to the Bloom’s syndrome disease gene. Mol Cell Biol, 18, 2721–2728.

30. Freyer, G.A., Davey, S., Ferrer, J.V., Martin, A.M., Beach, D. and Doetsch, P.W. (1995) An alternative eukaryotic DNA excision repair pathway. Mol Cell Biol, 15, 4572–4577.

31. Murray, J.M., Lindsay, H.D., Munday, C.A. and Carr, A.M. (1997) Role of Schizosaccharomyces pombe RecQ homolog, recombination, and checkpoint genes in UV damage tolerance. Mol Cell Biol, 17, 6868–6875.

32. Doe, C.L., Ahn, J.S., Dixon, J. and Whitby, M.C. (2002) Mus81-Eme1 and Rqh1 involvement in processing stalled and collapsed replication forks. J Biol Chem, 277, 32753–32759.

33. Caspari, T., Murray, J.M. and Carr, A.M. (2002) Cdc2-cyclin B kinase activity links Crb2 and Rqh1-topoisomerase III. Genes Dev, 16, 1195–1208.

34. Laursen, L.V., Ampatzidou, E., Andersen, A.H. and Murray, J.M. (2003) Role for the fission yeast RecQ helicase in DNA repair in G2. Mol Cell Biol, 23, 3692–3705.

35. Vindigni, A., Marino, F. and Gileadi, O. (2010) Probing the structural basis of RecQ helicase function. Biophys Chem, 149, 67–77.

36. Viziteu, E., Klein, B., Basbous, J., Lin, Y.L., Hirtz, C., Gourzones, C., Tiers, L., Bruyer, A., Vincent, L., Grandmougin, C. et al. (2017) RECQ1 helicase is involved in replication stress survival and drug resistance in multiple myeloma. Leukemia, 31, 2104–2113.

37. Sun, J., Wang, Y., Xia, Y., Xu, Y., Ouyang, T., Li, J., Wang, T., Fan, Z., Fan, T., Lin, B. et al. (2015) Mutations in RECQL Gene Are Associated with Predisposition to Breast Cancer. PLoS Genet, 11, e1005228.

38. Cybulski, C., Carrot-Zhang, J., Kluzniak, W., Rivera, B., Kashyap, A., Wokolorczyk, D., Giroux, S., Nadaf, J., Hamel, N., Zhang, S. et al. (2015) Germline RECQL mutations are associated with breast cancer susceptibility. Nat Genet, 47, 643–646.

39. Li, D., Frazier, M., Evans, D.B., Hess, K.R., Crane, C.H., Jiao, L. and Abbruzzese, J.L. (2006) Single nucleotide polymorphisms of RecQ1, RAD54L, and ATM genes are associated with reduced survival of pancreatic cancer. J Clin Oncol, 24, 1720–1728.

40. Zhi, L.Q., Ma, W., Zhang, H., Zeng, S.X. and Chen, B. (2014) Association of RECQL5 gene polymorphisms and osteosarcoma in a Chinese Han population. Tumour Biol, 35, 3255–3259.

41. Groocock, L.M., Prudden, J., Perry, J.J. and Boddy, M.N. (2012) The RecQ4 orthologue Hrq1 is critical for DNA interstrand cross-link repair and genome stability in fission yeast. Mol Cell Biol, 32, 276–287.

42. Mandell, J.G., Goodrich, K.J., Bahler, J. and Cech, T.R. (2005) Expression of a RecQ helicase homolog affects progression through crisis in fission yeast lacking telomerase. J Biol Chem, 280, 5249–5257.

43. Xu, Y.J. and Kelly, T.J. (2009) Autoinhibition and autoactivation of the DNA replication checkpoint kinase Cds1. J Biol Chem, 284, 16016–16027.

44. Xu, Y.J. (2016) Inner nuclear membrane protein Lem2 facilitates Rad3-mediated checkpoint signaling under replication stress induced by nucleotide depletion in fission yeast. Cell Signal, 28, 235–245.

45. Furuya, K., Poitelea, M., Guo, L., Caspari, T. and Carr, A.M. (2004) Chk1 activation requires Rad9 S/TQ-site phosphorylation to promote association with C-terminal BRCT domains of Rad4TOPBP1. Genes Dev, 18, 1154–1164.

46. Qu, M., Yang, B., Tao, L., Yates, J.R., 3rd, Russell, P., Dong, M.Q. and Du, L.L. (2012) Phosphorylation-dependent interactions between Crb2 and Chk1 are essential for DNA damage checkpoint. PLoS Genet, 8, e1002817.

47. Esashi, F. and Yanagida, M. (1999) Cdc2 phosphorylation of Crb2 is required for reestablishing cell cycle progression after the damage checkpoint. Mol Cell, 4, 167–174.

48. Capasso, H., Palermo, C., Wan, S., Rao, H., John, U.P., O’Connell, M.J. and Walworth, N.C. (2002) Phosphorylation activates Chk1 and is required for checkpoint-mediated cell cycle arrest. J Cell Sci, 115, 4555–4564.

49. Lopez-Girona, A., Tanaka, K., Chen, X.B., Baber, B.A., McGowan, C.H. and Russell, P. (2001) Serine-345 is required for Rad3-dependent phosphorylation and function of checkpoint kinase Chk1 in fission yeast. Proc Natl Acad Sci U S A, 98, 11289–11294.

50. Ivanova, T., Alves-Rodrigues, I., Gomez-Escoda, B., Dutta, C., DeCaprio, J.A., Rhind, N., Hidalgo, E. and Ayte, J. (2013) The DNA damage and the DNA replication checkpoints converge at the MBF transcription factor. Mol Biol Cell, 24, 3350–3357.

51. Dutta, C., Patel, P.K., Rosebrock, A., Oliva, A., Leatherwood, J. and Rhind, N. (2008) The DNA replication checkpoint directly regulates MBF-dependent G1/S transcription. Mol Cell Biol, 28, 5977–5985.

52. Lai, J.S. and Herr, W. (1992) Ethidium bromide provides a simple tool for identifying genuine DNA-independent protein associations. Proc Natl Acad Sci U S A, 89, 6958–6962.

53. Fernandez, A., Guo, H.S., Saenz, P., Simon-Buela, L., Gomez de Cedron, M. and Garcia, J.A. (1997) The motif V of plum pox potyvirus CI RNA helicase is involved in NTP hydrolysis and is essential for virus RNA replication. Nucleic Acids Res, 25, 4474–4480.

54. Moolenaar, G.F., Visse, R., Ortiz-Buysse, M., Goosen, N. and van de Putte, P. (1994) Helicase motifs V and VI of the Escherichia coli UvrB protein of the UvrABC endonuclease are essential for the formation of the preincision complex. J Mol Biol, 240, 294–307.

55. Muzzolini, L., Beuron, F., Patwardhan, A., Popuri, V., Cui, S., Niccolini, B., Rappas, M., Freemont, P.S. and Vindigni, A. (2007) Different quaternary structures of human RECQ1 are associated with its dual enzymatic activity. PLoS Biol, 5, e20.

56. Karow, J.K., Newman, R.H., Freemont, P.S. and Hickson, I.D. (1999) Oligomeric ring structure of the Bloom’s syndrome helicase. Curr Biol, 9, 597–600.

57. Compton, S.A., Tolun, G., Kamath-Loeb, A.S., Loeb, L.A. and Griffith, J.D. (2008) The Werner syndrome protein binds replication fork and holliday junction DNAs as an oligomer. J Biol Chem, 283, 24478–24483.

58. Hayles, J. and Nurse, P. (2018) Introduction to Fission Yeast as a Model System. Cold Spring Harb Protoc, 2018.

59. Lee, M.G. and Nurse, P. (1987) Complementation used to clone a human homologue of the fission yeast cell cycle control gene cdc2. Nature, 327, 31–35.

60. Shaag, A., Walsh, T., Renbaum, P., Kirchhoff, T., Nafa, K., Shiovitz, S., Mandell, J.B., Welcsh, P., Lee, M.K., Ellis, N. et al. (2005) Functional and genomic approaches reveal an ancient CHEK2 allele associated with breast cancer in the Ashkenazi Jewish population. Hum Mol Genet, 14, 555–563.

61. Yamagata, K., Kato, J., Shimamoto, A., Goto, M., Furuichi, Y. and Ikeda, H. (1998) Bloom’s and Werner’s syndrome genes suppress hyperrecombination in yeast sgs1 mutant: implication for genomic instability in human diseases. Proc Natl Acad Sci U S A, 95, 8733–8738.

62. Willis, N. and Rhind, N. (2009) Mus81, Rhp51(Rad51), and Rqh1 form an epistatic pathway required for the S-phase DNA damage checkpoint. Mol Biol Cell, 20, 819–833.

63. Marchetti, M.A., Weinberger, M., Murakami, Y., Burhans, W.C. and Huberman, J.A. (2006) Production of reactive oxygen species in response to replication stress and inappropriate mitosis in fission yeast. J Cell Sci, 119, 124–131.

64. Singh, A. and Xu, Y.J. (2016) The cell killing mechanisms of hydroxyurea. Genes (Basel), 7, 1–15.

65. Bjergbaek, L., Cobb, J.A., Tsai-Pflugfelder, M. and Gasser, S.M. (2005) Mechanistically distinct roles for Sgs1p in checkpoint activation and replication fork maintenance. EMBO J, 24, 405–417.

66. Durocher, D., Taylor, I.A., Sarbassova, D., Haire, L.F., Westcott, S.L., Jackson, S.P., Smerdon, S.J. and Yaffe, M.B. (2000) The molecular basis of FHA domain:phosphopeptide binding specificity and implications for phospho-dependent signaling mechanisms. Mol Cell, 6, 1169–1182.

67. Hishida, T., Han, Y.W., Shibata, T., Kubota, Y., Ishino, Y., Iwasaki, H. and Shinagawa, H. (2004) Role of the Escherichia coli RecQ DNA helicase in SOS signaling and genome stabilization at stalled replication forks. Genes Dev, 18, 1886–1897.

68. Parvathaneni, S. and Sharma, S. (2019) The DNA repair helicase RECQ1 has a checkpoint-dependent role in mediating DNA damage responses induced by gemcitabine. J Biol Chem, 294, 15330–15345.

69. Park, S.J., Lee, Y.J., Beck, B.D. and Lee, S.H. (2006) A positive involvement of RecQL4 in UV-induced S-phase arrest. DNA Cell Biol, 25, 696–703.

70. Park, S.Y., Kim, H., Im, J.S. and Lee, J.K. (2019) ATM activation is impaired in human cells defective in RecQL4 helicase activity. Biochem Biophys Res Commun, 509, 379–383.

71. Pichierri, P., Nicolai, S., Cignolo, L., Bignami, M. and Franchitto, A. (2012) The RAD9-RAD1-HUS1 (9.1.1) complex interacts with WRN and is crucial to regulate its response to replication fork stalling. Oncogene, 31, 2809–2823.

72. Patro, B.S., Frohlich, R., Bohr, V.A. and Stevnsner, T. (2011) WRN helicase regulates the ATR-CHK1-induced S-phase checkpoint pathway in response to topoisomerase-I-DNA covalent complexes. J Cell Sci, 124, 3967–3979.

73. Cheng, W.H., Muftic, D., Muftuoglu, M., Dawut, L., Morris, C., Helleday, T., Shiloh, Y. and Bohr, V.A. (2008) WRN is required for ATM activation and the S-phase checkpoint in response to interstrand cross-link-induced DNA double-strand breaks. Mol Biol Cell, 19, 3923–3933.

74. Davies, S.L., North, P.S., Dart, A., Lakin, N.D. and Hickson, I.D. (2004) Phosphorylation of the Bloom’s syndrome helicase and its role in recovery from S-phase arrest. Mol Cell Biol, 24, 1279–1291.

75. Moreno, S., Klar, A. and Nurse, P. (1991) Molecular genetic analysis of fission yeast Schizosaccharomyces pombe. Methods Enzymol, 194, 795–823.

76. Kowalczyk, K.M., Hartmuth, S., Perera, D., Stansfield, P. and Petersen, J. (2013) Control of Sty1 MAPK activity through stabilisation of the Pyp2 MAPK phosphatase. J Cell Sci, 126, 3324–3332.

